# Emerging mosquito resistance to piperonyl butoxide-synergized pyrethroid insecticide and its mechanism

**DOI:** 10.1101/2021.09.29.462303

**Authors:** Guofa Zhou, Yiji Li, Brook Jeang, Xiaoming Wang, Daibin Zhong, Guiyun Yan

**Affiliations:** Program in Public Health, University of California, Irvine, CA, USA

**Keywords:** *Culex quinquefasciatus*, insecticide resistance, piperonyl butoxide (PBO), PBO-ynergized pyrethroid, knockdown resistance, metabolic enzyme expressions

## Abstract

Piperonyl butoxide (PBO)-synergized pyrethroid products are widely available for the control of pyrethroid-resistant mosquitoes. To date, no study has formally examined mosquito resistance to PBO-synergized insecticides. We used *Culex quinquefasciatus* as a model mosquito examined the insecticide resistance mechanisms of mosquitoes to PBO-synergized pyrethroid using modified World Health Organization tube bioassays and biochemical analysis of metabolic enzyme expressions prior- and post-PBO exposure. We measured mosquito mortalities and metabolic enzyme expressions in mosquitoes with/without pre-exposure to different PBO concentrations and exposure durations. We found that field *Culex quinquefasciatus* mosquitoes were resistant to all insecticides tested, including PBO-synergized pyrethroids (mortality ranged from 3.7±4.7% to 66.7±7.7%), except malathion. Field mosquitoes had elevated levels of carboxylesterase (COE, 3.8-fold) and monooxygenase (P450, 2.1-fold) but not glutathione S-transferase (GST) compared to susceptible mosquitoes. When the field mosquitoes were pre-exposed to 4% PBO, the 50% lethal concentration of deltamethrin was reduced from 0.22% to 0.10%, compare to 0.02% for susceptible mosquitoes. Knockdown resistance gene mutation (L1014F) rate was 62% in field mosquitoes. PBO pre-exposure suppressed P450 enzyme expression levels by 25∼34%, GST by 11%, and had no impact on COE enzyme expression. Even with the optimal PBO concentration and exposure duration, field mosquitoes had significantly higher P450 enzyme expression levels after PBO exposure compared to laboratory controls. These results demonstrate that PBO alone may not be enough to control highly pyrethroid resistant mosquitoes due to the multiple resistance mechanisms. Mosquito resistance to PBO-synergized insecticide should be closely monitored.

**Authors’ Summary:** Mosquitoes are vectors of many major infectious diseases globally. Insecticides and related products are widely used for mosquito controls and disease preventions. Over time and following repeated use, mosquitoes (including *Aedes, Anopheles* and *Culex*) have developed very high resistance to multiple insecticides all over the world. Target site insensitivity due to mutations in the voltage-gated sodium channel gene and overproduction of metabolic detoxification enzymes such as cytochrome P450 (CYP) monooxygenases play critical role in insecticide resistance in mosquitoes. To enhance the killing power of insecticides, synergized insecticides were developed by mixing insecticide synergists with pyrethroids. Discovered in the 1940s, piperonyl butoxide (PBO) is one of the earliest and most commonly used insecticide synergists. Field application of PBO-synergized insecticides performed far better than mono-pyrethroids. PBO-treated long-lasting insecticidal nets (PBO-LLINs), which also use pyrethroids, outperformed regular LLIN for malaria control in many African countries. PBO-LLIN is soon to be rolled out on a large scale for malaria control in Africa. One important question regarding the use of synergized insecticides is whether they will select for vector population resistance to synergized insecticide products, in other words, are PBO-synergized pyrethroids effective against highly insecticide-resistant mosquitoes? To date, no study has formally examined mosquito resistance to PBO-synergized insecticides. Here, we used *Culex quinquefasciatus* as a model mosquito, we examined its resistance status to different insecticides including PBO-synergized pyrethrins and tested how PBO exposure affect mosquito mortality and the expressions of metabolic enzymes. We found that field *Culex quinquefasciatus* mosquitoes were resistant to multiple insecticides tested, including PBO-synergized pyrethroids. Field mosquitoes had elevated levels of carboxylesterase (COE) and monooxygenase (P450) but not glutathione S-transferase (GST) enzyme expressions compared to susceptible mosquitoes. Even with optimal PBO concentration and exposure duration, field mosquitoes had significantly higher P450 enzyme expression levels after PBO exposure compared to laboratory controls, and PBO exposure had no impact on COE enzyme expressions. The phenomena of the insecticide-resistant mosquitoes’ insensitivity to PBO exposure or PBO-synergized insecticides and multiple-resistance mechanisms have also been reported from *Aedes* and *Anopheles* mosquitoes in different countries. These results demonstrate that PBO alone is not enough to control highly pyrethroid resistant mosquitoes due to multiple resistance mechanisms. Mosquito resistance to PBO-synergized insecticide should be closely monitored

## 1. Introduction

Mosquitoes are well-known vectors of many infectious diseases including malaria, dengue, West Nile, yellow fever, Zika, chikungunya, and lymphatic filariasis, etc [1–3]. Among these diseases, dengue, chikungunya, Zika, yellow fever, Japanese encephalitis, non-*Plasmodium falciparum* malarias such as *Plasmodium vivax* and *Plasmosium knowlesi*, and lymphatic filariasis are considered as neglected tropical diseases [1]. Insecticide especially pyrethroid insecticide spraying and pyrethroid-treated bed nets are the most commonly used mosquito control methods [2,4,5]. Over time and following repeated use, mosquitoes have developed very high resistance to multiple insecticides all over the world [5–7]. A number of studies have examined mosquito pyrethroid resistance mechanisms and demonstrated that target site insensitivity due to mutations in the voltage-gated sodium channel (VGSC) gene, also known as knockdown resistance (*kdr*), has spread across the world [7]. The overproduction of metabolic detoxification enzymes such as cytochrome P450 (CYP) monooxygenases also plays critical role in insecticide resistance in mosquitoes [7–9].

To enhance the killing power of insecticides, synergized insecticides were developed by mixing insecticide synergists with pyrethroids [10–12]. The insecticide synergist by itself does not harm insects at low concentrations; instead, it enhances insecticides’ toxicity by inhibiting metabolic enzyme activities [10–12]. Discovered in the 1940s, piperonyl butoxide (PBO) is one of the earliest insecticide synergists [10–12]. PBO can synergize the effects of pyrethroid insecticides by reducing or nullifying the detoxifying capabilities of enzymes, primarily monooxygenases. PBO synergized pyrethoid ultra-low-volume sprayings have been widely used for *Aedes* and *Culex* mosquito controls [13–16]. Field application of PBO-synergized insecticides performed far better than mono-pyrethroids [10,12]. In addition to PBO, S,S,S-tributyl phosphorotrithioate (DEF), triphenyl phosphate (TPP), and diethyl maleate (DEM) are used as synergists for other classes of insecticides [17–19]. However, PBO is the most popularly used insecticide synergist [16,17]. Today, more than 2,500 EPA-registered pesticide products in the USA contain PBO synergist [20]. PBO-treated long-lasting insecticidal nets (PBO-LLINs), which also use pyrethroids, have been tested for malaria control in many African countries [21], with results showing that PBO-LLIN outperformed regular LLIN in reducing malaria infections and vector densities [21,22]. PBO-LLIN is soon to be rolled out on a large scale for malaria control in Africa with the support of the President’s Malaria Initiatives [23].

One important question regarding the use of synergized insecticides is whether they will select for vector population resistance to synergized insecticides; in other words, are PBO-synergized pyrethroids effective against highly insecticide-resistant mosquitoes? Laboratory selection studies found that the addition of PBO slowed the development of pyrethroid insecticide resistance in *Aedes aegypti* and *Anopheles stephensi* mosquitoes and agricultural insect pest white flies; however, insecticide-resistant mosquitoes quickly developed resistance to PBO-synergized insecticides [24–26]. A study in *Musca domestica* houseflies found that the flies developed resistance to PBO-synergized pyrethrins after just one year of continuous field applications of PBO-synergized pyrethrins [27]. A recent study in Mozambique found that highly insecticide-resistant *Anopheles funestus* was also resistant to PBO-LLIN [28]. A number of other studies also found that pre-treatment of insecticide-resistant *Aedes, Anopheles* and *Culex* mosquitoes did not fully restore the mosquitoes’ susceptibility to insecticides and sometimes had a very limited effect on mortality [29–31]. There are several possible causes for mosquito to tolerant PBO-synergized insecticides, including: A) the PBO concentration is too low or the exposure duration is too short; B) the resistance is so intense that PBO exposure cannot overcome the limit of the resistance, i.e., there is a maximum limit to the effect of PBO; C) mosquitoes develop tolerance to the PBO synergist, which is unlikely in places where PBO products have not been used, however, it is unknown in places such as in the USA where PBO synergized insecticide products have been used for some time [16]; and D) other resistance mechanisms are present, e.g., multiple mechanisms. However, no study has concurrently extensively examined mosquito resistance status to PBO-synergized pyrethroid insecticide resistance and its mechanisms.

The aim of this study was to use *Cx. quinquefasciatus* as a model mosquito to examine levels of mosquito resistance to PBO-synergized pyrethroids and the underlying mechanisms in mosquitoes collected from Southern California, where PBO-synergized pyrethroids have not been used. We tested PBO exposure against both susceptible and field-collected mosquitoes and examined *kdr* mutations and metabolic resistance mechanisms. The results will be useful for guiding the large-scale use of PBO-synergized insecticide products, facilitating development of new methods of surveilling insecticide resistance, and informing the development of new vector control strategies.

## 2. Materials and Methods

### 2.1 Ethics statement

No permits were required for the described field mosquito collections. Mosquito collections in breeding sites were consented to orally by the property owners at each location. This study did not involve the collection of any human-related samples or personal information such as participants’ names, addresses, and GPS location of their homes.

### 2.2 *Culex quinquefasciatus* populations and colonies

Field *Cx. quinquefasciatus* (OC-R strain) larvae were collected from multiple places in Irvine, Garden Grove and Orange cities of Orange County, Southern California. *Cx. quinquefasciatus* in Southern California are resistant to pyrethroids through elevated metabolic detoxification and knockdown resistance mechanisms [6,32,33]. Notably, PBO-synergized pyrethroid insecticides have not been applied outdoors in field settings in Orange County and surrounding areas. Thus, we presume that the field collected *Culex* used in our study had never been exposed to PBO-synergized pyrethroids. The eggs of an insecticide-susceptible *Cx. quinquefasciatus* strain, JHB F179, were obtained from BEI Resources (BEI Resources, NIAID, NIH, MRA No. NR-43025). All mosquitoes were reared in insectaries in the Program in Public Health, University of California, Irvine. The conditions of mosquito rearing and at which all resistance experiments were conducted in this study are as follows: 26 ± 1°C, relative humidity 70 ± 10%, and light : dark ratio of 12 h : 12 h. F1 females of the OC-R strain were used in the study, and F0 females of the JHB F179 strain were used. All emerged adults were maintained with cotton balls soaked in 10% sugar solution before any experiments.

### 2.3 Test of mosquito insecticide resistance

We used the World Health Organization (WHO) standard tube bioassay to determine the resistance of *Cx. quinquefasciatus* mosquitoes to different insecticides [34]. We tested four insecticides: the pyrethroids, 1) deltamethrin (concentration 0.05%, 0.25% and 0.5%) and 2) permethrin (0.75%); 3) the carbamate, bendiocarb (0.1%); and 4) the organophosphate, malathion (5%). Silicone oil was used as the control. Females 3–5 days old were exposed to insecticide-impregnated papers for 1 hour, then transferred to a holding tube and provided with cotton balls soaked in 10% sugar solution. Knockdown was recorded every 10 min. Mortality was determined after 24 h. The field OC-R strain (F1) was tested against all four insecticides; for JHB F179 (F0), only the pyrethroids were tested. Twenty females were used for each test, with 4 replicates of treatment and 2 replicates of control performed for each insecticide.

### 2.4 Mosquito resistance to PBO-synergized pyrethroid

We used USA-marketed PBO pyrethrin as the reference insecticide. We used a mix of 6% pyrethrin and 60% PBO plus silicone oil. We impregnated Whatman No. 1 (1 mm) paper using a solution for indoor contact and surface spray, i.e., 4.25 fl oz/gallon of water and 1 gallon/750 square feet, or approximately 0.947 ml of solution per test paper (12 cm × 15 cm). We further diluted the solution to 1.5 ml per test paper before impregnating test paper. To examine the mortality of JHB F179 and OC-R females, we used PBO-synergized pyrethrin impregnated test paper and the WHO tube bioassay as described above with slight modifications, i.e., 20 females per test, 2 h exposure and mortality scored after 24 h, with 4 replicates for treatment and 2 for control (silicone oil test paper).

### 2.5 PBO effect – extended exposure time and increased concentration

For the first set of tests, we used a modified version of WHO-recommended PBO synergist-insecticide bioassays [34], i.e., PBO 4% and deltamethrin 0.05%. Twenty 3–5-day-old females were exposed to PBO-impregnated papers for 1 h, followed by 1 h exposure to insecticide-impregnated papers. Mosquitoes were then transferred to a holding tube and provided with cotton balls soaked in 10% sugar solution. Knockdown was recorded every 10 min. Mortality was determined after 24 h. We additionally tested PBO exposure times of 2, 3, 4, and 6 hours. Twenty females were used for each test, with 4 replicates for PBO + deltamethrin and 2 replicates for each control (PBO- and silicone oil-impregnated papers). The field OC-R strain (F1) was tested for all treatments; for JHB F179 (F0), only the exposure condition of PBO 1h + deltamethrin 1h was tested.

To determine the optimal PBO concentration and exposure time, we tested deltamethrin 0.05% alone and with pre-exposure to PBO at concentrations of 4%, 7%, and 10%, with exposure times of 1 h, 2 h, and 3 h using field OC-R strain mosquitoes.

### 2.6 Resistance intensity: Dose-response relationship and synergistic effect

To determine the dose-response relationship between PBO synergist exposure and mosquito susceptibility to pyrethroids, we conducted WHO synergist-insecticide bioassays using different insecticide concentrations. The deltamethrin concentrations tested are listed in Appendix Table S1. PBO concentration was 4%, which is currently the WHO-recommended diagnostic concentration. The test procedures were the same as described above, i.e., four replicates for treatment and two replicates for control, duration of exposure was 1 h with PBO followed by 1 h with insecticide. Mosquito mortality was scored after 24 h and recorded for all tests.

### 2.7 Knockdown resistance and metabolic enzyme activity measurement

Identification of L1014F *kdr* mutations was conducted on untreated mosquitoes using a TaqMan assay based on Yoshimizu et al. [32]. Among the field *Culex*, only three mosquitoes were found dead 24 h after deltamethrin exposure. Therefore, kdr mutations were examined using pooled mosquitoes.

Cytochrome P450 monooxygenase enzyme activity was examined for four populations: the susceptible JHB F179 strain without exposure to deltamethrin insecticide or PBO, the field OC-R strain before and after 1 h exposure to 4% PBO (WHO standards), and OC-R after 3 h exposure to 7% PBO (which yielded robust mortality from this study). We also examined activities of glutathione S-transferases (GSTs) and carboxylesterases (COEs). We modified a previously published protocol to measure monooxygenase and GST activities [35]. Details of the protocols are described in Appendix File 1. Total protein was measured for each mosquito and corrected against a bovine serum albumin (BSA Fraction V; Sigma) standard curve. All measurements were done in triplicate. Mean absorbance values for each tested mosquito and enzyme were converted into enzyme activity and standardized based on the total protein amount.

### 2.8 Data analysis

WHO susceptibility test criteria are as follows: susceptible if mortality rate ≥ 98%; possible resistance if mortality is 90–97%; confirmed resistance if mortality rate < 90% or mortality rate < 98% with repeated tests using untested mosquitoes from the population exhibiting possible resistance as described prior; moderate to high intensity resistance if mortality < 98% with 5× the diagnostic concentration; and high intensity resistance if mortality < 98% with 10× the diagnostic concentration. We used *χ*^2^-tests to determine the differences in mortalities between tests with different insecticides, different insecticide concentrations of the same insecticide, different PBO exposure durations, different PBO concentrations, and the same insecticide with/without PBO pre-exposure. A *χ*^2^-test was also used to determine the difference in mortalities between field and susceptible mosquitoes exposed to PBO-synergized pyrethrin after 2 h exposure. Lethal concentration was estimated using the Probit model. LC50 and LC90, concentrations that respectively killed 50% and 90% of mosquitoes, were estimated for susceptible and field strains of mosquitoes with/without PBO pre-exposure. Resistance ratio, RR, was calculated as (LC50 or LC90 of field mosquitoes)/(LC50 or LC90 of susceptible mosquitoes). PBO synergistic ratio, SR, was calculated as (LC50 or LC90 of field mosquitoes with PBO)/(LC50 or LC90 of field mosquitoes without PBO).

The difference in *kdr* rates between susceptible and field mosquitoes was not analyzed because susceptible mosquitoes did not show any *kdr* mutations. Analysis of variance (ANOVA) was used to compare P450, GST, and COE enzyme levels between different mosquito strains without exposure to insecticide or PBO, and between field mosquitoes before and after exposure to different concentrations of PBO. Reduction in enzyme activities due to PBO exposure was calculated by comparing enzyme levels before and after PBO exposure.

## 3. Results

### 3.1 Intensity of insecticide resistance in *Cx. quinquefasciatus*

WHO tube bioassays revealed that using the diagnostic dosages, mortality of field *Cx. quinquefasciatus* was <10% against pyrethroids (i.e., deltamethrin and permethrin), 66.7±7.7% against bendiocarb, and 96.3±4.7% against malathion, while susceptible JHB F179 mosquitoes were not fully susceptible to deltamethrin (97.5±1.7%) (Figure 1A). Even with 5× and 10× increased deltamethrin concentrations, field *Cx. quinquefasciatus* mortalities were <60% (Figure 1A). For the field mosquitoes, knockdown started late with standard diagnostic insecticide dosages, and the knockdown rate was low by the end of 1 h exposure. The knockdown rate increased significantly with 5× and 10× deltamethrin (Figure 1B).

**Figure 1.**
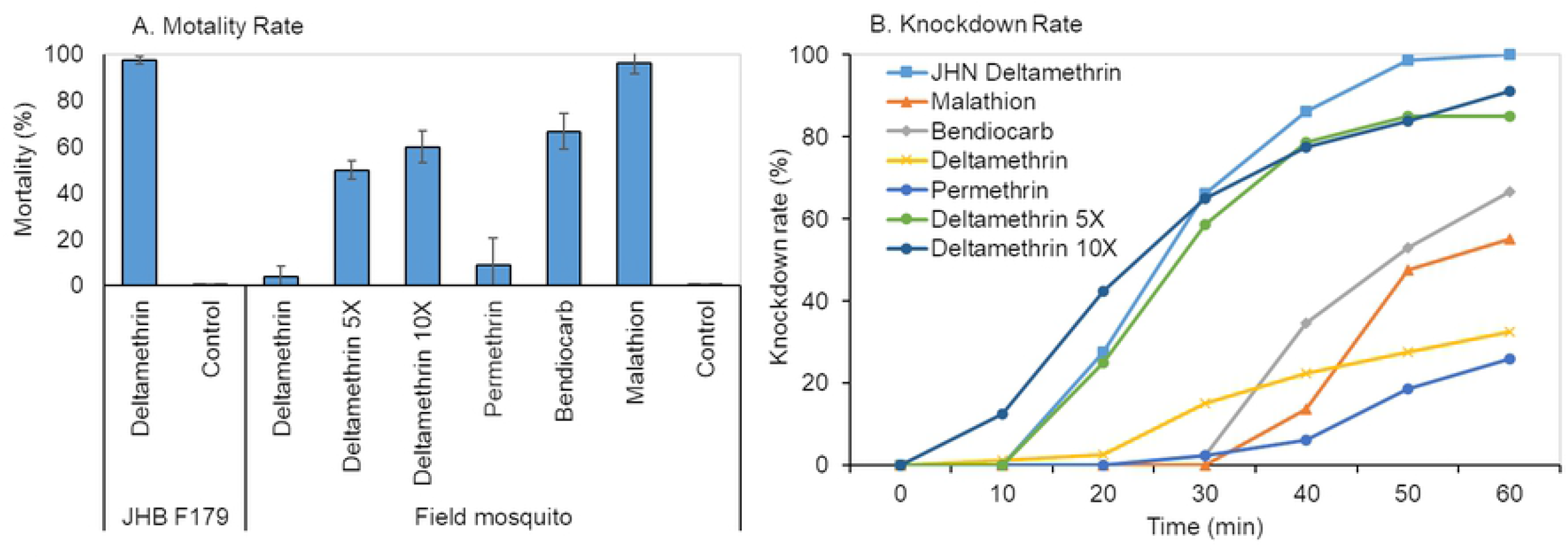
Mortality of female *Cx. quinquefasciatus* mosquitoes against different insecticides based on WHO tube test using standard diagnostic concentrations and 5× and 10× deltamethrin concentrations. A) Mortality rate (%); and B) Knockdown rate (%). JHB deltamethrin refers to JHB F179 strain mosquitoes exposed to deltamethrin; all others curves represent field strain mosquitoes and the insecticides they were exposed to. As controls, field and JHB F179 strain mosquitoes were separately exposed to silicone oil impregnated paper.

### 3.2 *Cx. quinquefasciatus* against PBO-synergized pyrethroid

After 2 h exposure of *Cx. quinquefasciatus* to PBO-synergized pyrethrins (1:10 pyrethrins : PBO by volume), the mortality rate was 91.3% (95% CI: 84.0–98.6%) and the knockdown rate 98.8% (95% CI: 96.0–100%) for the JHB F179 colony, whereas the field population had a mortality rate of 1.6% (95% CI: 0–4.4%) and a knockdown rate of 13.3% (95% CI: 5.8–20.8%) (Figure 2). Both the mortality and knockdown rates were significantly higher among JHB F179 strain mosquitoes compared to field mosquitoes (*χ*^2^-test, P < 0.01).

**Figure 2.**
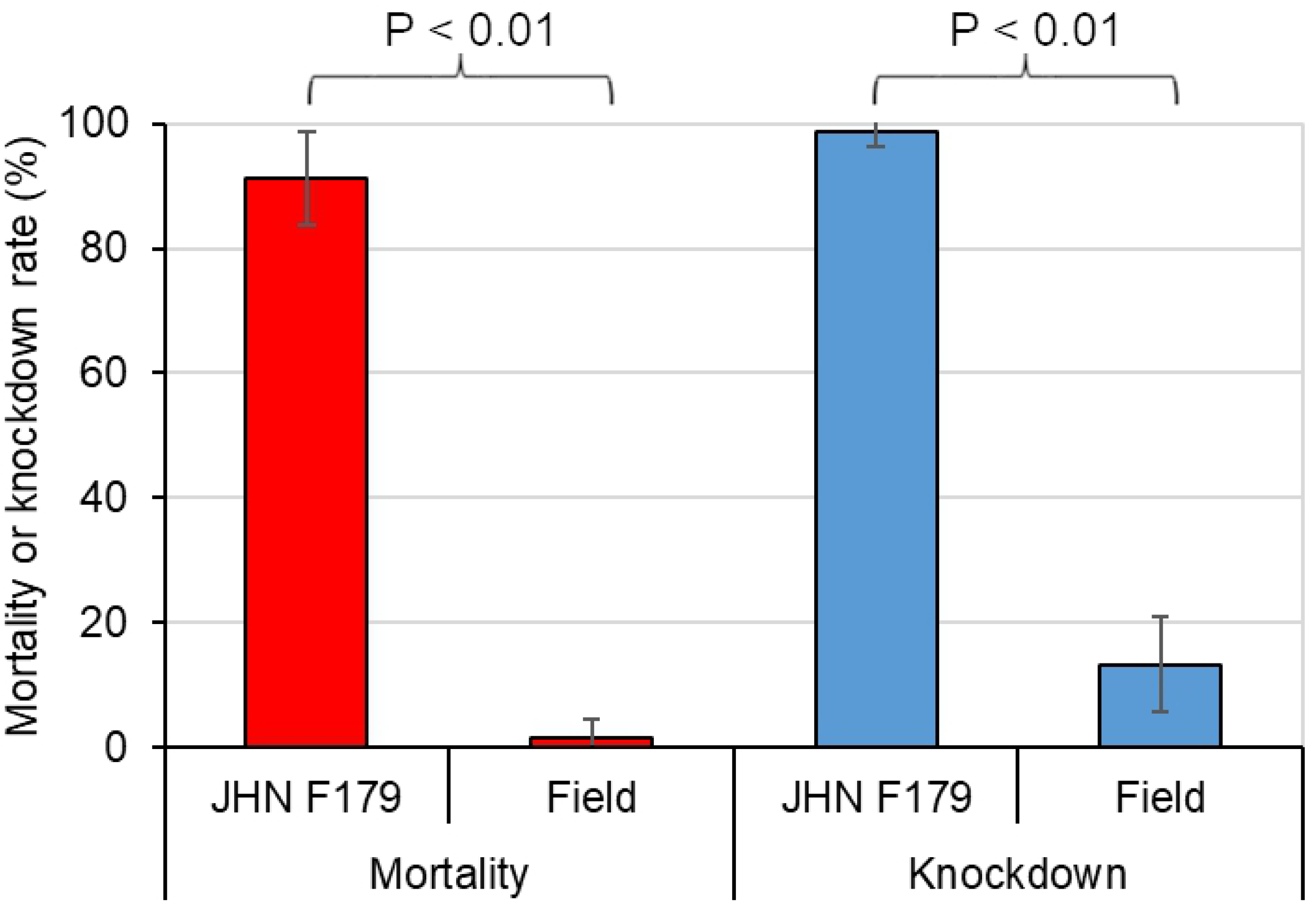
Mortality and knockdown rates of female mosquitoes against PBO-synergized pyrethrin after 2 h exposure.

### 3.3 PBO effect on mosquito mortality

For OC-R mosquitoes, pre-exposure to 4% PBO for 1 h more than doubled mosquito mortality; however, mortality rates remained <10% even after PBO pre-exposure (Figure 3A). Extending the duration of 4% PBO exposure to 2–6 h increased mosquito mortality against deltamethrin; however, mortality remained < 30% (Figure 3A), and mortalities were similar between exposure durations from 2–6 h (Figure 3A).

**Figure 3.**
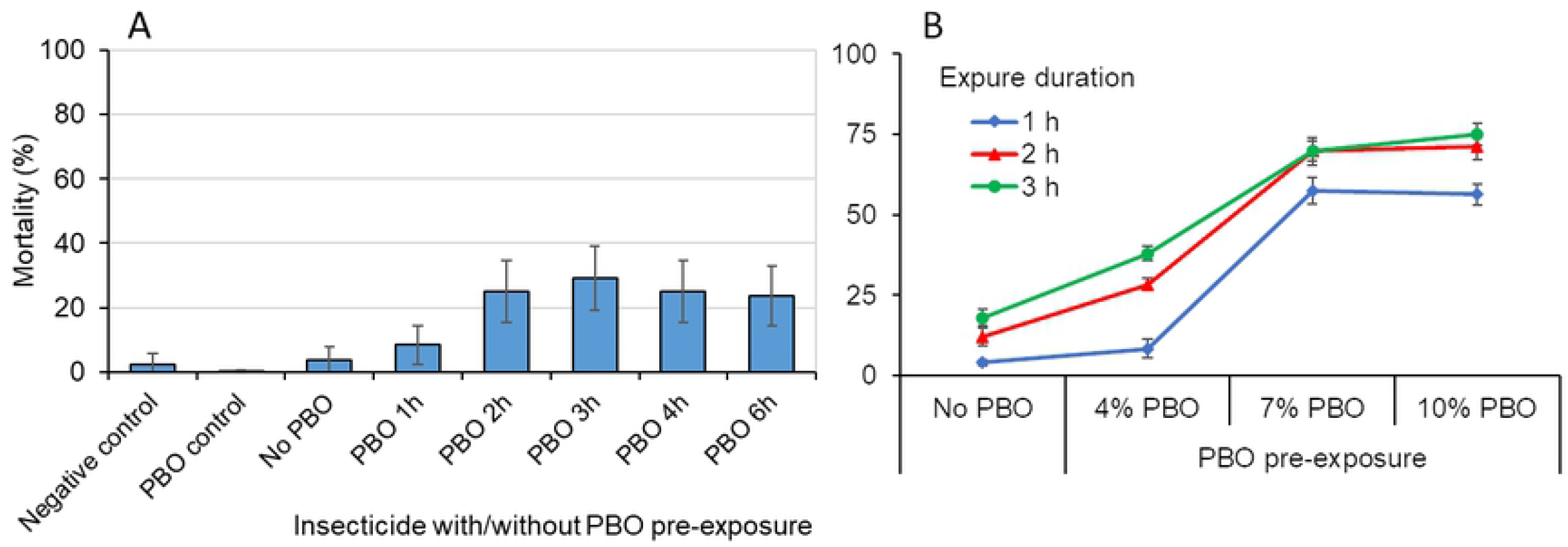
Mortality (%) of field *Cx. quinquefasciatus* against different PBO concentrations and exposure durations. A) No PBO and no insecticide (negative control), PBO control (no insecticide), and insecticide with 0–6 h pre-exposure to 4% PBO; and B) 1–3 h pre-exposure to 4–10% PBO.

Bioassays with 0.05% deltamethrin without PBO pre-exposure resulted in mortality rates <20% after 1–3 hours of exposure, confirming high pyrethroid resistance in the field *Cx. quinquefasciatus* mosquitoes (Figure 3B). PBO pre-exposure led to higher mortality against deltamethrin for PBO concentrations up to 7%. Mortality at 10% PBO concentration was very similar to that at 7% (Figure 3B). Increasing the PBO exposure duration from 1 h to 2 h led to higher mortality rates, but no significant difference in mortality was observed between 2 h and 3 h PBO exposure (P > 0.05, Figure 3B).

### 3.4 Dose-response relationship and PBO synergistic ratio

Dose-response relationships for the JHB F179 and OC-R mosquitoes against deltamethrin with/without PBO are shown in Figure 4. The 50% lethal concentration (LC50) of the OC-R population was 0.22% without PBO pre-exposure, indicating a resistance ratio of 11.2 in relation to the susceptible JHB F179 colony (Table 1). When the mosquitoes were pre-exposed to 4% PBO for 1 h, LC50 was reduced to 0.10%, and the resistance ratio decreased to 4.9 with a synergistic ratio of 2.3 (Table 1). Similar resistance and synergistic ratios were observed using LC90 (Table 1).

**Table 1.** Estimation of 50% and 90% lethal concentrations of deltamethrin, resistance ratio (RR), and PBO synergistic ratio (SR).

| LC <sup>†</sup> | Mosquito strain | Concentration (%) |  | RR <sup>‡</sup> | SR <sup>§</sup> |
| --- | --- | --- | --- | --- | --- |
|  |  | No PBO | With PBO |  |  |
| LC50 <sup>†</sup> | Susceptible strain | 0.0200 | - | - | - |
|  | Field strain | 0.2246 | 0.0989 | 11.2 | 2.3 |
| LC90 <sup>†</sup> | Susceptible strain | 0.0248 | - | - | - |
|  | Field strain | 0.3028 | 0.1351 | 12.2 | 2.2 |

<sup>†</sup> LC: lethal concentration; LC50: LC that kills 50% of mosquitoes; LC90: LC that kills 90% of mosquitoes; ‘-’: not applicable.
<sup>‡</sup> RR = (LC50 or LC90 of susceptible strain)/(LC50 or LC90 of field strain)
<sup>§</sup> SR = (LC50 or LC90 of field strain with no PBO)/(LC50 or LC90 of field strain with PBO).

**Figure 4.**
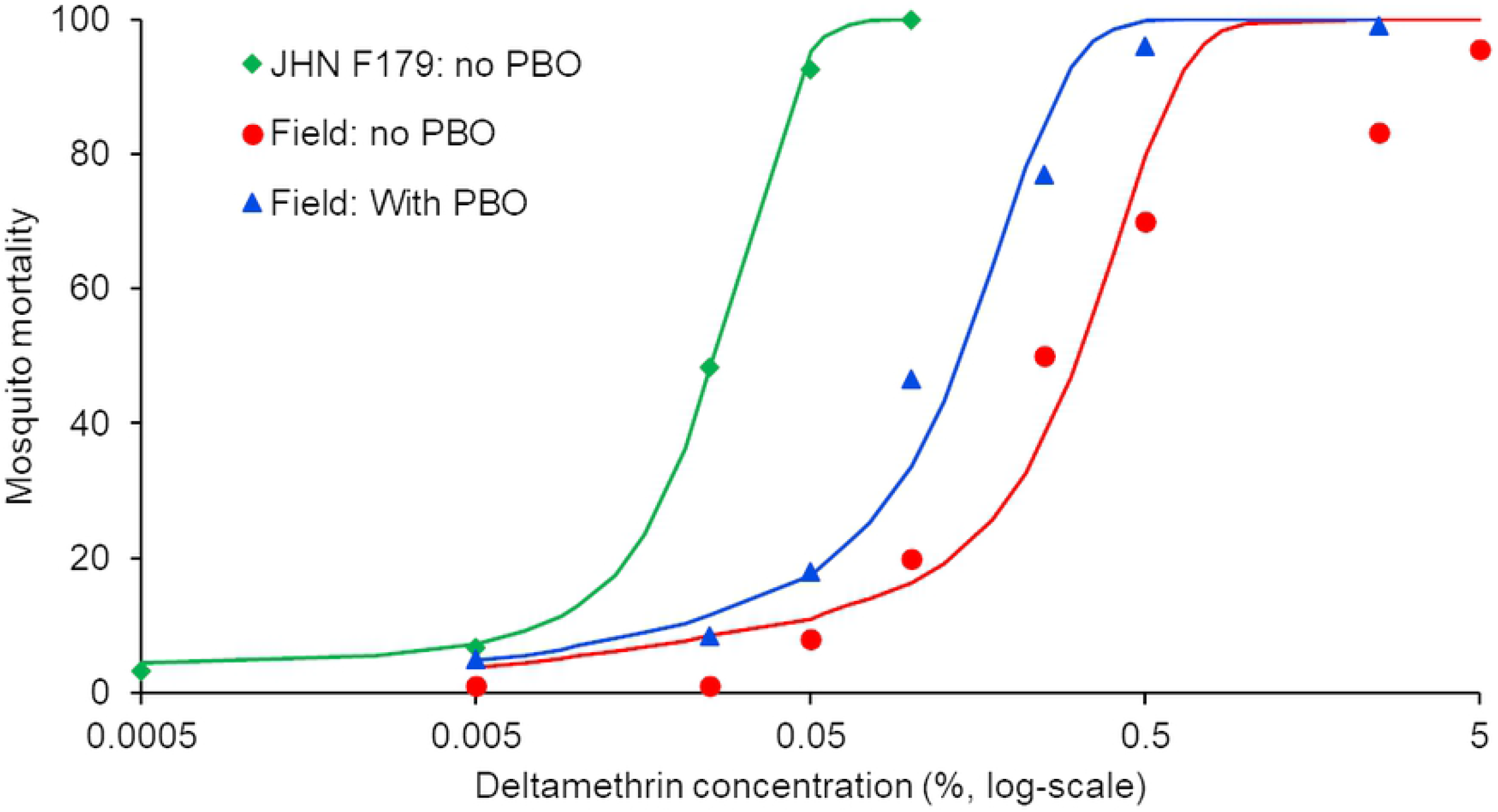
Dose-response relationship against deltamethrin and PBO exposure. Dots/triangles/diamonds represent observed mortalities; curves represent model fitted results. Green (diamond), blue (triangle), and red (dot) colors respectively represent JHB F179 without PBO pre-exposure, field mosquitoes without PBO pre-exposure, and field mosquitoes with PBO pre-exposure.

### 3.5 Knockdown gene mutations and metabolic enzyme expressions

The knockdown resistance mutation rate for L1014 was 61.8% in OC-R mosquitoes without exposure to pyrethroid insecticides and 0% in JHB F179 mosquitoes.

Compared to the JHB F179 susceptible mosquitoes, field mosquitoes had significantly elevated enzyme activity for P450 (2.1-fold) and COE (3.8-fold) but not for GST (1.0-fold) (ANOVA, P < 0.05) (Figure 5, Table 2 and Additional Table S2). For the field mosquitoes, pre-exposure to 4% PBO reduced P450 enzyme levels by about 25% but did not affect COE or GST enzyme activity. Pre-exposure to 7% PBO significantly suppressed enzyme activity for both P450 (33%) and GST (11%) but not for COE (ANOVA, P < 0.05) (Figure 5, Table 2 and Additional Table S2). Even after 7% PBO exposure, P450 enzyme levels in field mosquitoes were still significantly higher than enzyme levels in JHB F179 mosquitoes (1.4-fold, ANOVA, P < 0.05).

**Figure 5.**
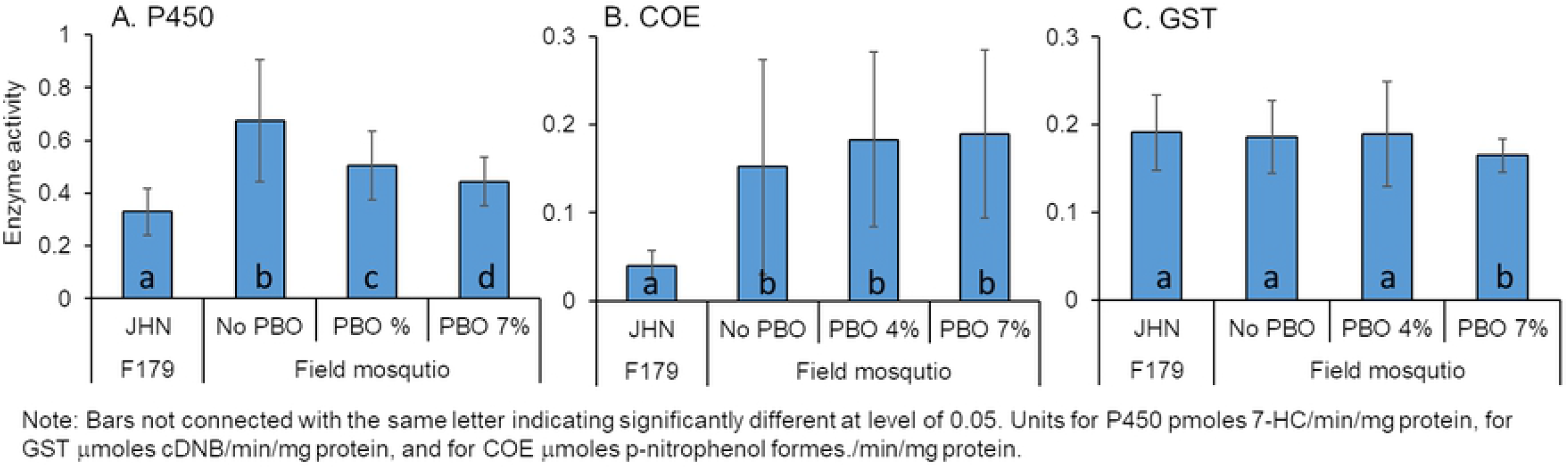
Standardized enzyme activities based on the total protein amount for different mosquito strains and for field mosquitoes at different PBO concentrations.

**Table 2.** Ratio of enzyme levels between field and susceptible mosquitoes and between different PBO treatments of field mosquitoes.

| Enzyme | Field control<br>vs. JHB F179 | PBO 4% exposure<br>vs. field control | PBO 7% exposure<br>vs. field control |
| --- | --- | --- | --- |
| GST † | 0.97 | 1.02 | 0.89 ** |
| COE † | 3.80 *** | 1.20 | 1.24 |
| P450 † | 2.05 *** | 0.75 *** | 0.66 *** |

† GST: glutathione S-transferase; COE: carboxylesterase; P450: monooxygenase.
\*\* Significantly different at level of 0.01; \*\*\* significantly different at level of 0.0001;
otherwise, not significantly different at level of <0.05.

## 4. Discussion

Mosquito resistance to insecticides is a major challenge for mosquito-borne disease prevention and control. PBO-synergized pyrethroid insecticides are one way to combat insecticide resistance. However, PBO-synergized insecticide products have their own limitations. Although PBO-synergized pyrethroids are widely used in the USA for *Aedes* and *Culex* controls [13–16], and PBO-LLINs have been tested and are rolling out in many malaria endemic African countries [21–23], potential mosquito resistance to PBO-synergized pyrethroids and PBO-LLIN has not attracted attention from the scientific community and policy makers. Here, we used *Cx. quinquefasciatus* as a model mosquito to examine the interactions between highly insecticide-resistant mosquitoes and PBO exposure. We found that field *Cx. quinquefasciatus* from Southern California were highly resistant to pyrethroid and carbamate insecticides. Although PBO-synergized pyrethroids have never been used for outdoor mosquito control in Orange County, CA, field *Cx. quinquefasciatus* were highly resistant to market formulation equivalent PBO-synergized pyrethroids. In addition, even with optimal PBO concentration and exposure duration, PBO pre-exposure paired with the diagnostic dose of deltamethrin killed only about 70% of field *Cx. quinquefasciatus*, indicating the limitation of PBO exposure on mosquito mortality. Biochemical analysis revealed that PBO could significantly reduce P450 enzyme activity; however, the reduced P450 activity in field *Cx. quinquefasciatus* was still significantly higher than that in the susceptible mosquitoes. This may partly explain why PBO pre-exposure and PBO-synergized pyrethroids could not fully restore mosquitoes’ susceptibility to insecticides. There was highly elevated COE enzyme activity in field mosquitoes, and exposure to PBO had no impact on COE enzyme activity. In other words, the impact of PBO-synergized pyrethroids on mosquito mortality is limited because of multiple mechanisms, so other synergists such as DEF are essential to counteract the mosquitoes’ resistance to insecticides.

The phenomena of the insecticide-resistant mosquitoes’ insensitivity to PBO exposure or PBO-LLIN have also been reported for *Aedes* and *Anopheles* mosquitoes from different countries [28,31,36–40]. For example, in Marcombe and colleagues’ study in Laos, for highly permethrin-resistant *Ae. aegypti*, exposure to PBO only increased mosquitoes’ mortalities from 2-11% to 10-25% [31]. Although one may argue that mortalities have increased 2-5 folds, however, the observed mortalities are way too low from mosquito control point of view. In Singapore, Koou et al reported very similar findings for *Ae. aegypti* mosquitoes against cypermethrin, permethrin and etofenprox [37]. Similarly, for *An. gambiae* s.l. study in Mali, Keita and colleagues’ found that in several places PBO exposure could only increase mosquito mortalities to pyrethroid insecticides from 24-41% to 34-45% [40]. In Riveron et al study of *An. funestus* in Mozambique, PBO exposure increased mosquito mortalities to pyrethroids from about 10% to <30%; in addition, mosquito mortalities increased from around 5% against regular LLIN to about 15% against PBO-LLIN even after 72 h of exposure [28]. We have noted that PBO-synergized insecticide products have not been used in above mentioned study areas. These results suggest that highly insecticide-resistant mosquitoes may tolerant PBO exposure regardless of species. In nearly all cases, multiple resistance mechanisms have been reported, including presence of different detoxifying enzymes and different *kdr* mutations [37–40].

Although PBO-synergized pyrethrum and pyrethroid insecticides have been tested and used in the field since 1950 [12,27,41], insect resistance to PBO-synergized insecticides and insecticide products is seldom reported [27,28]. Davies *et al*. first reported housefly resistance to PBO pyrethrins in 1958, and Riveron *et al*. reported *Anopheles funestus* resistance to PBO-pyrethroid treated long-lasting insecticidal nets in 2019 [27,28]. Many studies found that pre-exposure of highly pyrethroid-resistant *Aedes, Anopheles* and *Culex* mosquitoes to a synergist such as PBO, DEF, TPP or DEM did not fully restore the mosquitoes’ mortality [36,42–47], however, most of these studies used the synergist to infer the mechanisms behind resistance without examining the mechanisms themselves, i.e., enzyme activity before and after synergist exposure. For example, most studies using PBO 4% + insecticide found that PBO pre-exposure increased the mortality of field mosquitoes [28,42,43,48]; thus, researchers concluded that P450 enzyme was involved in resistance. To the best of our knowledge, our study is the first to examine in detail both enzyme activities and bioassay tests after mosquitoes were exposed to PBO. We examined how PBO exposure, at different concentrations and exposure durations, affected mosquito mortality as well as P450, COE, and GST enzyme activities.

Although increased enzyme activity and expression in P450 CYP genes have been well recognized as the major mechanisms for pyrethroid resistance [8,40,42,43,49,50], it is unclear why the effect of PBO exposure is limited in reducing resistance levels. One possibility is the existence of multiple mechanisms of resistance. It is generally believed that PBO affects mainly P450 enzyme activity [51–53]. Other studies found that both mixed function oxidase and esterase activity were involved in *Ae. aegypti* and *Cx. quinquefasciatus* resistance to PBO-synergized insecticides, while resistant *Anopheles* also developed GST and A296S-RDL dieldrin resistance mechanisms [31,37,42,49,50,54]. In this study, we found that both P450 and COE enzyme activities in field *Cx. quinquefasciatus* were significantly elevated compared to those in mosquitoes of the susceptible strain, whereas GST levels were similar between the susceptible and field mosquitoes. This is similar to what has been found in cases of extremely high and multiple insecticide resistance in the malaria mosquito *Anopheles gambiae*, i.e., gene mutations in P450 enzymes and acetylcholinesterase ACE-1 duplication in COE enzymes [53]. We find about 60% *kdr* gene mutations in field *Culex* mosquitoes, which is also in line with previous study conducted in the same area [32]. More interestingly, in addition to multiple resistance mechanisms, we found that even with robust PBO concentration and exposure duration, the reduction in P450 enzyme activity in field mosquitoes was limited, this is a complete new finding. Rather than multiple mechanisms of resistance, we suspect that PBO has a maximum carrying capacity with regard to reducing P450 enzyme activity. When mosquito resistance exceeds a certain level, PBO exposure cannot overcome such a high level of enzyme activity; i.e., the synergistic effect of PBO has a limit, however, need more evidences. We cannot rule out other unknown mechanisms.

Another currently available PBO product is the PBO-LLIN. The chief reason for producing PBO-LLIN is the widespread high pyrethroid resistance levels in *Anopheles* mosquitoes in malaria-endemic African countries. Although the majority of previous studies found that PBO-LLIN outperformed LLIN at increasing killing power and/or reducing malaria infections [21–23,55], several studies reported reduced efficacy of PBO-LLIN against highly resistant *Anopheles* and *Culex* mosquitoes [28,30,49,53–55]. We should note that it is unlikely that African *Anopheles* mosquitoes were exposed to PBO or PBO-synergized insecticide products before large-scale rollout of PBO-LLINs. As discussed earlier, the reduced PBO-LLIN killing efficacy is likely due to multiple resistance mechanisms [49,50,53,54]; i.e., adding PBO to LLIN may not be enough to combat *Anopheles* mosquitoes’ resistance to pyrethroids.

This study has some limitations. First, WHO insecticide susceptibility test guidelines do not include standard diagnostic insecticide doses for *Culex* mosquitoes [34]. We used the standard diagnostic insecticide doses for *Anopheles* mosquitoes in this study [34]. We tested 5X and 10X diagnostic insecticide doses and observed that the mortalities of field collected *Culex* mosquitoes were all <60%. In addition, mortality of JHB F179 control strain *Culex* was 97% against standard 1X diagnostic insecticide doses. These results collectively suggest that field-collected *Culex* mosquitoes were highly resistant to pyrethroid insecticides. Second, we acknowledge that commonly applied control methods for *Culex* and *Aedes* mosquito management are aerial/ground ultra-low-volume (ULV) adulticiding and larviciding [5,56]. ULV spraying of PBO-synergized pyrethroids such as Aqualuer® 20-20 has been extensively evaluated [56,57]. Although semi-field (caged) experiments using direct spraying can be adapted to evaluate the efficacy of PBO-synergized pyrethroids [57,58], the outcomes are affected by many factors including the specifications of sprayers, travel distance of droplets, and environmental conditions (e.g., wind speed and direction) [56–60], which likely contributed to inter-experimental variations in mosquito mortality [56]. Therefore, we did not conduct direct spraying experiments. Last but not least, results from laboratory-controlled tests likely differ from results following actual field applications [56]. However, results are comparable under the same experimental conditions. Field applications of ULV spraying show large variations in reductions in mosquito densities [56]; it is unknown whether the low effectiveness is due to mosquitoes’ tolerance to insecticides, spraying methods, or the coverage of spraying, as it is infeasible for aerial/ground sprayings to cover every mosquito [56,60]. Furthermore, field environmental conditions also affect spraying efficacy [56]. Therefore, laboratory-controlled environment tests are a viable alternative for the evaluation of the efficacy of PBO-synergized insecticides. Indeed, we observed 92% mortality of JHB 179 control strain *Culex* mosquitoes against PBO-synergized pyrethrins compared to 2% mortality among our field mosquitoes. This indicates validity of our laboratory-controlled tests as well as field *Culex* tolerance of PBO-synergized pyrethrins.

## 5. Conclusion

*Cx. quinquefasciatus* from Southern California were highly resistant to PBO-synergized pyrethroids where PBO-synergized pyrethroids has never been used outside in the field. This raises serious concerns about the efficacy of currently available PBO synergized insecticide products against mosquitoes, including PBO-synergized pyrethroids for spraying and for LLINs. With the increased use of insecticides with or without PBO, mosquitoes will develop increased resistance to insecticides, which will potentially compromise the efficacy of PBO-synergized insecticide products. Potential development of mosquito tolerance to PBO-synergized insecticide products should be closely monitored.

## Acknowledgments

We wish to thank the Orange County Mosquito and Vector Control District, CA, for their support in collecting some of the field mosquitoes used in this study. This study was partially funded by the National Institutes of Health (D43 TW01505).

## Support Information

Appendix tables

**Table S1.** Deltamethrin concentrations tested for different mosquito strains and PBO treatments.

| Deltamethrin<br>concentration (%) | JHB F179 | Field mosquito |  |
| --- | --- | --- | --- |
|  | No PBO | No PBO | 4% PBO |
| 0.0005 | √ |  |  |
| 0.005 | √ | √ | √ |
| 0.025 | √ | √ | √ |
| 0.05 | √ | √ | √ |
| 0.1 | √ | √ | √ |
| 0.25 |  | √ | √ |
| 0.5 |  | √ | √ |
| 2.5 |  | √ |  |
| 5 |  | √ |  |

**Table S2.** Standardized enzyme expression (mean ± SD) based on the total protein amount and standard curve

| Enzyme <sup>†</sup> | JHB F179 | Field mosquito |  |  |
| --- | --- | --- | --- | --- |
|  |  | Control | PBO 4% exposure | PBO 7% exposure |
| GST <sup>‡</sup> | 0.191 ± 0.043 a | 0.186 ± 0.041 a | 0.189 ± 0.060 a | 0.165 ± 0.019 b |
| COE <sup>‡</sup> | 0.040 ± 0.017 a | 0.152 ± 0.122 b | 0.183 ± 0.099 c | 0.189 ± 0.095 c |
| P450 <sup>‡</sup> | 0.329 ± 0.088 a | 0.674 ± 0.232 b | 0.503 ± 0.131 c | 0.443 ± 0.092 d |

<sup>†</sup> Comparisons were done using Tukey HSD test; different lowercase letters in the same row indicate significant difference at a level of 0.05.
<sup>‡</sup> GST: glutathione S-transferases, $\mu$ moles/min/mg protein; COE: carboxylesterases, $\mu$ moles/min/mg protein; P450: monooxygenase, mg cytochrome C equivalents/mg protein.

